# *In vitro* efficacy of synthetic antimicrobial peptide SET-M33 against poultry isolates with diverse antimicrobial resistance phenotypes

**DOI:** 10.64898/2026.05.12.724496

**Authors:** Ana Laura Pereira Lourenço, Alessia Maranesi, Gerardo Ceada, Teresa Ayats, Nuria Aloy, Nuria Navarro, Noelia Antilles, Mar Biarnés, Chiara Falciani, Alessandro Pini, Karl Kochanowski, Marta Cerda Cuellar

**Affiliations:** Unitat Mixta d’Investigació IRTA UAB en Sanitat Animal, Centre de Recerca en Sanitat Animal (CReSA), Campus de la Universitat Autònoma de Barcelona (UAB), 08193 Bellaterra, Barcelona, Spain; IRTA, Programa de Sanitat Animal, Centre de Recerca en Sanitat Animal (CReSA), Campus de La UAB, 08193 Bellaterra (Cerdanyola del Vallès), Spain; Department of Material Science and Engineering, Universitat Politècnica de Catalunya, Barcelona, 08019, Spain; Department of Cell Biology, Physiology and Immunology, Universitat Autònoma de Barcelona, Bellaterra, 08193, Spain; Centre de Sanitat Avícola de Catalunya i Aragó(CESAC), 43206 Reus (Tarragona), Spain; University of Siena. Department of Medical Biotechnology, 53100 Siena, Italy; Clinical Pathology Unit, Azienda Ospedaliera Universitaria Senese, via Mario Bracci, 53100 Siena, Italy; SetLance srl, 53100 Siena, Italy

**Keywords:** Antimicrobial peptides, Antimicrobial resistance, Chicken intestinal organoids, Poultry, Multidrug resistance

## Abstract

Antimicrobial resistance is an impactful One Health issue. One of its drivers is the extensive use of antibiotics in both human and animal production systems, and despite regulatory restrictions on antibiotic use in poultry farming, antimicrobial resistance remains a major challenge. Consequently, animals are at higher risk of harder-to-treat diseases and play a role as resistance reservoirs, highlighting the need for alternative antimicrobial strategies. Towards this end, antimicrobial peptides (AMPs) have emerged as promising candidates due to their broad-spectrum activity and lower propensity to induce resistance. However, the effectiveness of AMPs against poultry pathogens, and in particular multi drug-resistant strains, is largely unclear. To tackle this question, we evaluated the synthetic AMP SET-M33 against four species of clinically relevant pathogens in poultry, namely *Escherichia coli, Salmonella enterica, Enterococcus faecalis* and *Enterococcus cecorum*. Using a panel of 141 field isolates, we found that SET-M33 broadly inhibited bacterial growth at low micromolar concentrations (median MICs of 2.5 μM and 5 μM for Gram-negative and Gram-positive strains, respectively), including in multi drug-resistant isolates. To examine the potential impact of SET-M33 on the host, we established a new *in vitro* co-cultivation system using chicken intestinal organoids. We found that SET-M33 retains its antimicrobial activity in organoid-microbe co-cultures at concentrations that preserved host viability. These findings demonstrate the potential of SET-M33 as a new antimicrobial agent against pathogens in poultry.

## Introduction

Antimicrobial resistance (AMR) is a natural outcome of microbial evolution. However, the extensive and often inappropriate use of antibiotics in human medicine and food animal production has dramatically accelerated the emergence and dissemination of resistant bacteria, posing a major global health threat (1,2). AMR is widely recognized as a One Health issue, as it simultaneously affects human, animal, and environmental health through interconnected pathways (3,4). In food-producing animals, antibiotic use can select for resistant bacterial populations that may be transmitted to humans via the food chain or disseminated into the environment through animal waste, and vice versa (5,6). Once released, resistance genes can spread among bacterial communities through horizontal gene transfer, facilitating their reintroduction into both animal and human populations via agriculture and water systems (7). This self-sustaining cycle underscores both the complexity of AMR and the difficulty of reversing resistance once it becomes established.

Within this context, poultry production systems represent a critical interface in the AMR cycle (8,9). Resistant bacteria remain prevalent in poultry farms, despite the European Union banning the use of antibiotics for growth promotion in animal production in 2006 (10), and the entry into force of Regulation (EU) 2019/6 on veterinary medicinal products in 2022 (11) which introduced strict rules on antibiotic use to combat AMR. Numerous studies report that commensal and zoonotic bacteria isolated from both conventional and antibiotic-free systems frequently exhibit resistance to multiple antimicrobial classes (5,12–14). The persistence of resistance even in the absence of direct antibiotic selective pressure is likely driven by the low fitness cost associated with certain resistance mechanisms, particularly those acquired through horizontal gene transfer (15). These observations indicate that regulatory measures and reductions in antibiotic use, while essential, are insufficient on their own to control AMR in poultry production and highlight the need for novel antimicrobial agents capable of overcoming existing resistance determinants.

The development of such agents requires alternative modes of action that differ fundamentally from those of conventional antibiotics, and antimicrobial peptides (AMPs) have emerged as promising candidates (16,17). AMPs are naturally occurring components of the innate immune system across all domains of life and can also be synthetically engineered to enhance stability, activity, and safety (18–26). Their mechanisms of action are often multimodal and include interactions with bacterial membranes and other non-specific targets (27), enabling broad-spectrum activity against both susceptible and multidrug-resistant pathogens (28–32). Importantly, AMPs generally exhibit a lower propensity for inducing resistance (33–35), although resistance development cannot be entirely excluded (36,37).

In poultry production, several bacterial species are of particular relevance due to their clinical impact and their role as antimicrobial resistance vectors (8,38). *Escherichia coli* is a major cause of avian colibacillosis, leading to substantial economic losses and serving as a reservoir of resistance genes with implications for human health (38–41). Whilst birds normally act as asymptomatic carriers of *Salmonella* spp, infections associated with acute and chronic diseases in avian species may occur (42). However, the attention to *Salmonella* is largely due to its role as one of the most important foodborne pathogens worldwide, which is frequently associated with poultry products and shows high prevalence of multidrug-resistance despite the fact that birds are not treated with antibiotics to combat this pathogen (43). Therefore, AMPs may represent a valuable tool to reduce or eradicate *Salmonella* in poultry and thus prevent its entry into the food chain. *Enterococcus* spp., although commonly present as commensals in the avian gut, have also been associated with diseases in poultry and humans (44,45). Enterococci are relevant in AMR context due to their intrinsic and acquired resistance to several antibiotics (38,46). Together, these organisms represent priority targets for evaluating alternative antimicrobial strategies in poultry production systems.

In this study, we aimed to investigate the synthetic AMP SET-M33 as a potential alternative antimicrobial against the clinically relevant poultry pathogens *E. coli, Salmonella enterica, Enterococcus faecalis* and *Enterococcus cecorum*. SET-M33 was originally identified through a random phage display library against *E. coli* and subsequently optimized into a tetra-branched structure to enhance its antimicrobial properties (47). Previous studies have demonstrated the therapeutic potential of SET-M33 *in vitro* and *in vivo* in infection models relevant to human health (47–50). However, its efficacyagainst poultry-associated bacterial pathogens, particularly those exhibiting resistance to conventional antimicrobials, and its potential impact on the host itself, are currently unclear.

Here, we tackled these questions *in vitro* using a two-pronged approach: first, we determined the *in vitro* activity of SET-M33 against a large panel of 141 field strains of *E. coli, S. enterica, E. faecalis*, and *E. cecorum*. Our data revealed that SET-M33 broadly inhibits *in vitro* growth of these isolates at low micromolar concentrations, including in isolates that are resistant against multiple classes of conventional antibiotics. Second, we determined the antimicrobial activity of SET-M33 in presence of host cells using a new chicken intestinal organoid infection model. We found that SET-M33 effectively inhibited pathogen growth of most tested isolates at concentrations that did not impair host viability. Thus, this work highlights the potential of AMPs for tackling antimicrobial resistance in poultry production and developing broader One Health–oriented mitigation strategies.

## Methods

### Reagents and bacterial strains

Unless stated otherwise, all reagents were purchased from Sigma-Aldrich. Peptide SET-M33 was provided by SetLance (Italy). Avian field strains of *E. coli, S. enterica* ser. Enteritidis, Infantis and Typhimurium, *E. faecalis* and *E. cecorum* used in the study (**Figure 1A**) originated from northeastern Spain and were kindly supplied by Centre de Sanitat Avícola de Catalunya i Aragó(CESAC). *E. coli* strains included avian pathogenic *E. coli* (APEC) and non-APEC, according to the presence of 5 virulence genes (51).

**Figure 1.**
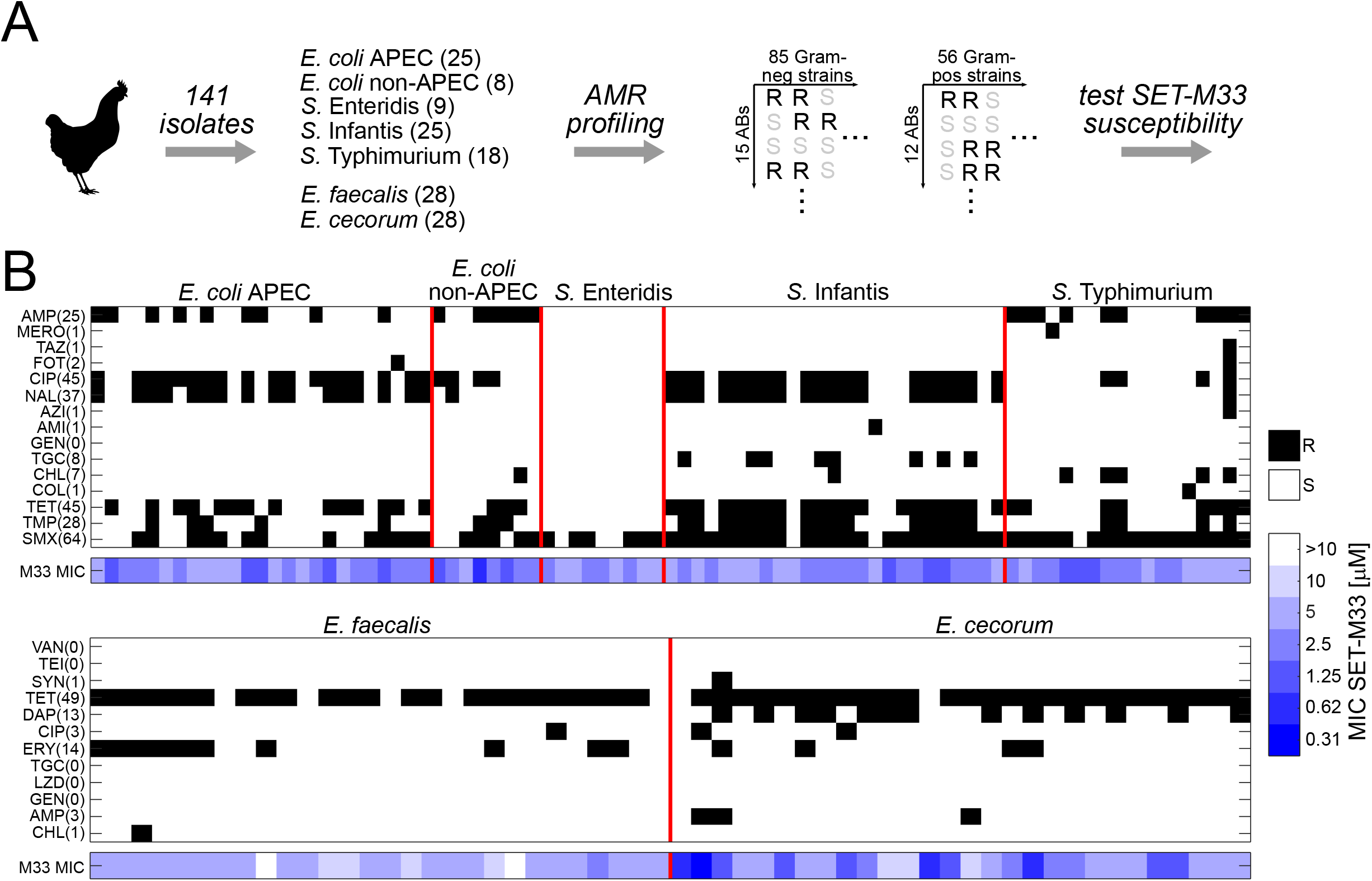
Antimicrobial resistance (AMR) profiling of poultry strains. **A)** Schematic of approach **B)** Overview of antibiotic resistance profiles for 85 Gram-positive strains (top row) tested against 15 antibiotics, and 56 Gram-positive strains (bottom row) tested against 12 antibiotics. Black squares: resistant (as determined following EUCAST guidelines). Numbers next to each antibiotic: number of resistant strains detected. Lower part: corresponding SET-M33 minimal inhibitory concentration (MIC). APEC, avian pathogenic E. coli. Antibiotics: ampicillin (AMP), meropenem (MERO), ceftazidime (TAZ), cefotaxime (FOT), ciprofloxacin (CIP), nalidixic acid (NAL), azithromycin (AZI), amikacin (AMI), gentamicin (GEN), tigecycline (TGC), chloramphenicol (CHL), colistin (COL), tetracycline (TET), trimethoprim (TMP), sulfamethoxazole (SMX), vancomycin (VAN), teicoplanin (TEI), quinupristin/dalfopristin (SYN), daptomycin (DAP), erythromycin (ERY), linezolid (LZN).

### Antimicrobial susceptibility testing

All field strains were tested for antibiotic susceptibility against antibiotics included in the EU surveillance program of AMR in food-producing animals by determining the minimum inhibitory concentration (MIC) using the commercial EUVSEC3 (for *E. coli* and *Salmonella*) and EUVENC (for *Enterococcus* species) microdilution plates (Thermo Fisher Scientific) according to the manufacturer’s instructions and EUCAST guidelines, except for *E. cecorum* in which methodology followed Laurentie et al. (52). Obtained MIC values were used to classify isolates as susceptible or resistant based on epidemiological cutoff (ECOFF) according to EUCAST (www.eucast.org) and Laurentie et al. (52). Isolates were considered to be multidrug-resistant (MDR) if they presented resistance to at least three classes of antibiotics (53). The full list of antibiotics used here is shown in **Supplementary Table 1**.

Apart from conventional antibiotics, isolates were also screened against the AMP SET-M33 with an in-house microdilution assay as described previously (54). Briefly, SET-M33 was tested in cation adjusted Mueller-Hinton broth (CAMHB, Thermofisher Scientific) (for *E. cecorum* CAMHB supplemented with 5%lysed horse blood was used) using different final concentrations ranging from 0.31 to 10 µM in 2-fold dilutions. Bacterial suspensions were prepared from fresh overnight Columbia blood agar (bioMérieux) cultures in PBS to McFarland 0.5 and used to inoculate 96-well plates (Greiner Cat. No 655 180) to a final concentration of 5 x 10^5^ CFU/mL, in a final volume of 100 µL per well. After incubation at 37 °C for 24h, results were collected both by visual inspection, checking absence/presence of growth, and by OD600 measurements using an automated plate reader (Tecan Nano M+, Tecan) for confirmation.

### Chicken intestinal organoid culture and passaging

A chicken jejunum organoid culture from a 23-days-old broiler chicken (Ross 308) was obtained from the organoid biobank at IRTA-CReSA (https://www.irta.cat/en/servei/zoorganoids-platform/). A vial of cryopreserved organoids was thawed, diluted in PBS (21-040-CV, Corning) and centrifuged at 300 x g for 5 min at 4°C. The pellet was resuspended in Matrigel®(45356231, Corning) and plated in small drops of ∼10uL in a 24-well plate (4-5 drops per well). The plate was incubated upside-down for 30 min at 37°C, 5%CO_2_, 95%humidity for the Matrigel®to solidify. Finally, the Matrigel®domes were overlayed with 500 µL/well of Human IntestiCult™organoid growth medium (OGM, 06010, Stemcell Technologies) supplemented with 1/100 Penicillin-Streptomycin (15140-122, Gibco) and 1/500 Amphotericin B (15290-026, Gibco) and incubated at 37°C, 5%CO_2_, 95%humidity for up to one week. Organoid growth was monitored using phase - contrast light microscopy and the medium was replaced every 3–4 days.

After one week, the organoids were passaged as follows: the 3D organoids were harvested from the wells by vigorous pipetting and collected into a 15-ml tube. Ice-cold PBS was used to recover residual organoids in the well and was collected into the same tube. The tube was centrifuged at 300 ×g for 5 min at 4°C and the supernatant was discarded. The organoids were dissociated by adding 1 ml of TrypLE™Express (12605036, ThermoFisher Scientific) to the pellet and incubating for 3 min at 37°C. The TryPLE was diluted with 4mL of PBS, and the tube was centrifuged at 300 ×g for 5 min at 4°C. The supernatant was discarded and the pellet was resuspended in Matrigel®to seed small drops in a 24-well plate, as indicated before. After 30 min of incubation upside-down at 37°C, 5%CO_2_, 95%humidity, 500 µl of OGM were dispensed in each well.

### Generation of organoid-derived 2D monolayers

A 96 well-plate was coated with a solution of rat-tail collagen I in PBS (10 µg/cm^2^) for 1h at 37°C. The collagen solution was then aspirated, and the wells were washed with PBS. 3D organoids were harvested from the wells by vigorous pipetting and collected in a 15-ml tube. Ice-cold PBS was used to recover residual organoids in the well and were collected into the same tube. Organoids were pelleted by centrifugation at 300 ×g for 5 min at 4 °C. The supernatant was discarded and the organoids were dissociated by adding 1 ml of TrypLE™Express followed by incubation for 7 min at 37°C. To ensure appropriate dissociation, the cell suspension was pipetted vigorously. TrypLE™Express was inactivated by adding 4 ml of PBS. The cells were centrifuged once again at 300 x g for 5 min at 4°C, and the supernatant was discarded. The pellet was resuspended in 500 µl of OGM and the cell concentration and viability were estimated via trypan blue exclusion with an automatic cell counter. Cell density was adjusted to seed 5 ×10^4^–6 ×10^4^ alive cells per well in the collagen-coated 96-well plate (100µL/well) in OGM. The organoid-derived 2D monolayer reached confluency after approximately 3 days of culture under standard incubation conditions (37°C, 5%CO2, 95%of humidity), as determined visually using phase-contrast light microscopy.

### Quantifying SET-M33 toxicity in organoid-derived 2D monolayers

Once organoid-derived 2D monolayers reached confluency, the spent media was discarded, and advanced (ADV) DMEM/F-12 supplemented with varying concentrations of SET-M33 was added (0.5-4 µM). The organoids were incubated for 24h at 37°C, 5%CO_2,_ 95%of humidity. Following incubation, the metabolic activity of the cells was quantified with CellTiterGlo®assay (G7570, Promega Corporation) according to the provider protocol. Briefly, CellTiterGlo®reagent was thawed at room temperature and 100 µl were added in each well. Wells without cells but with CellTiterGlo®served as background. The plate was shaken at 200 rpm for 2 min at room temperature (RT) and then incubated for 10 min statically at RT. The luminescence was measured in each well using a luminescence microplate reader (Fluoroskan FL, Thermo Fisher Scientific). Viability was calculated as the relative luminescence value compared to the negative control (untreated organoid-derived 2D monolayers in ADV DMEM/F-12) after background subtraction.

### Effect of SET-M33 on epithelial barrier integrity

The effect of the peptide on the epithelial barrier integrity of chicken intestinal organoid derived monolayers was assessed as follows: Organoid derived monolayers were seeded in 24 well cell culture inserts (0.4 µm PET clear extended, 9320422, cellQart) following the same procedure described in section “Generation of organoid-derived 2D monolayers”. The monolayers were incubated at 37°C, 5%CO_2,_ 95%of humidity and the epithelial barrier integrity was monitored by measuring trans epithelial electric resistance (TEER) with Epithelial Volt-Ohm Meter (Millicell ERS-2, MERS00002, Merck Millipore). Once the culture was confluent, the OGM media was discarded, the monolayers were washed with ADV DMEM/F-12 and 100 µl of SET-M33L solution at 3 µM were added to each well. The TEER was measured at different time points (0,2,4,6,24h) to investigate the effect of the peptide on the epithelial layer. Wells with only ADV DMEM /F-12 were used as controls.

### Antimicrobial activity of SET-M33 in chicken intestinal organoid infection model

Confluent organoid-derived 2D monolayers were infected with field strains of *E. coli* (15845, 13027-1, 15632), *S*. Enteritidis (8830-1, 19804-1, 25373-1) *S*. Infantis (12902-1, 14473-1, 26458-1) and *S*. Typhimurium (13089-1, 2352-1, 23721-1), to evaluate the antimicrobial efficacy of SET-M33. For each bacterial strain, colonies were collected from freshly prepared cultures on Columbia blood agar plates and suspended in PBS to obtain a 0.5 McFarland turbidity. The bacterial suspension was subsequently diluted in ADV DMEM/F-12 to achieve a final concentration of 1.5 ×10^6^ CFU/mL. Prior to infection, OGM was removed from the wells, and monolayers were gently washed with 100 µL of ADV DMEM/F12 to remove residual growth factors and antibiotics. SET-M33 was prepared in ADV DMEM/F12 and added to the monolayers at concentrations corresponding to 1×MIC, ∼1.5×MIC, and 2×MIC for each tested pathogen. Wells without SET-M33 were used as controls. For bacterial strains with a MIC of 2.5 µM, the maximum concentration tested was limited to 1.5×MIC to not exceed the previously established cytocompatibility range. A 50 µL of the peptide solution was added to each well, followed by 50 µL of the prepared bacterial suspension, resulting in co-incubation of monolayers with both pathogen and antimicrobial peptide. Infected cultures were incubated for 6 h at 37°C, 5%CO2, 95%of humidity. Following incubation, bacterial growth was quantified by serial dilution of culture supernatants in PBS and plating onto Columbia blood agar plates. Bacterial count was performed after overnight incubation at 37°C to assess antimicrobial efficacy.

### Data processing

Experimental data were processed in Excel (Microsoft) and custom MATLAB scripts (Mathworks, version R2021a).

## Results

To better understand the potential use of antimicrobial peptide SET-M33 to combat avian and zoonotic pathogens, we focused on four of the most important pathogen species. Specifically, we selected two Gram-negative species, namely *E. coli* and *Salmonella* spp., and two Gram-positive ones, namely *E. faecalis* and *E. cecorum*. From these four species, we obtained 141 field strains recovered during 2024 and 2025 from diseased animals, except for *Salmonella* which were collected within the framework of the poultry surveillance program. Isolates were selected from the strain collection as representative of different clinical cases and different poultry batches. To elucidate the prevalence of antibiotic resistances within this strain collection, we determined MICs against panels of conventional antibiotics –15 against Gram-negative and 12 against Gram-positive isolates, respectively (full list in **Supplementary Table 1**). In parallel, we determined the MIC for SET-M33 using an in-house microdilution assay with varying concentrations of 0.31 to 10 µM, matching the reported effective *in vivo* concentration range for this peptide (55,56) (see **Figure 1A** for a schematic of the approach).

The data revealed that the vast majority of strains (i.e. 78/85 and 49/56 for Gram-positive and Gram-negative strains) were resistant against at least one of the tested antibiotics (**Figure 1B**). We did observe some recurring resistance patterns. For example, most Gram-negative strains were resistant to the tested quinolones ciprofloxacin and nalidixic acid (i.e. 66.7%and 51.5%resistant strains, respectively), consistent with EFSA surveillance data in *E. coli* strains isolated from broilers (57). Similarly, most tested Gram-positive strains were resistant to tetracycline, again consistent with recent surveillance data (57). However, the data also revealed substantial differences in the number of resistances across strains, with as many as ten resistances found in individual strains. In comparison, susceptibility to SET-M33 tended to be more similar across the tested strains and, generally, MICs were in the low micromolar range. SET-M33 MIC values tended to be higher in Gram-positive strains, especially for *E. faecalis*, consistent with observations in Gram-positive pathogens (58). Nevertheless, only two out of 141 strains were able to grow at the highest tested SET-M33 concentration (10 µM).

Next, we explored the relationship between antibiotic resistance patterns and SET-M33 susceptibility in more detail. Specifically, we focused on multi-drug resistant (MDR) strains (see **Figure 2A** for a schematic of the approach). With the exception of S. ser. Infantis, all tested species had at least some MDR isolates (**Supplementary Figure 1**). When comparing SET-M33 susceptibility in MDR and non-MDR strains, we found no differences in the distribution of SET-M33 MICs, neither for Gram-negative (**Figure 2B**) nor for Gram-positive (**Figure 2C**) isolates.

**Figure 2.**
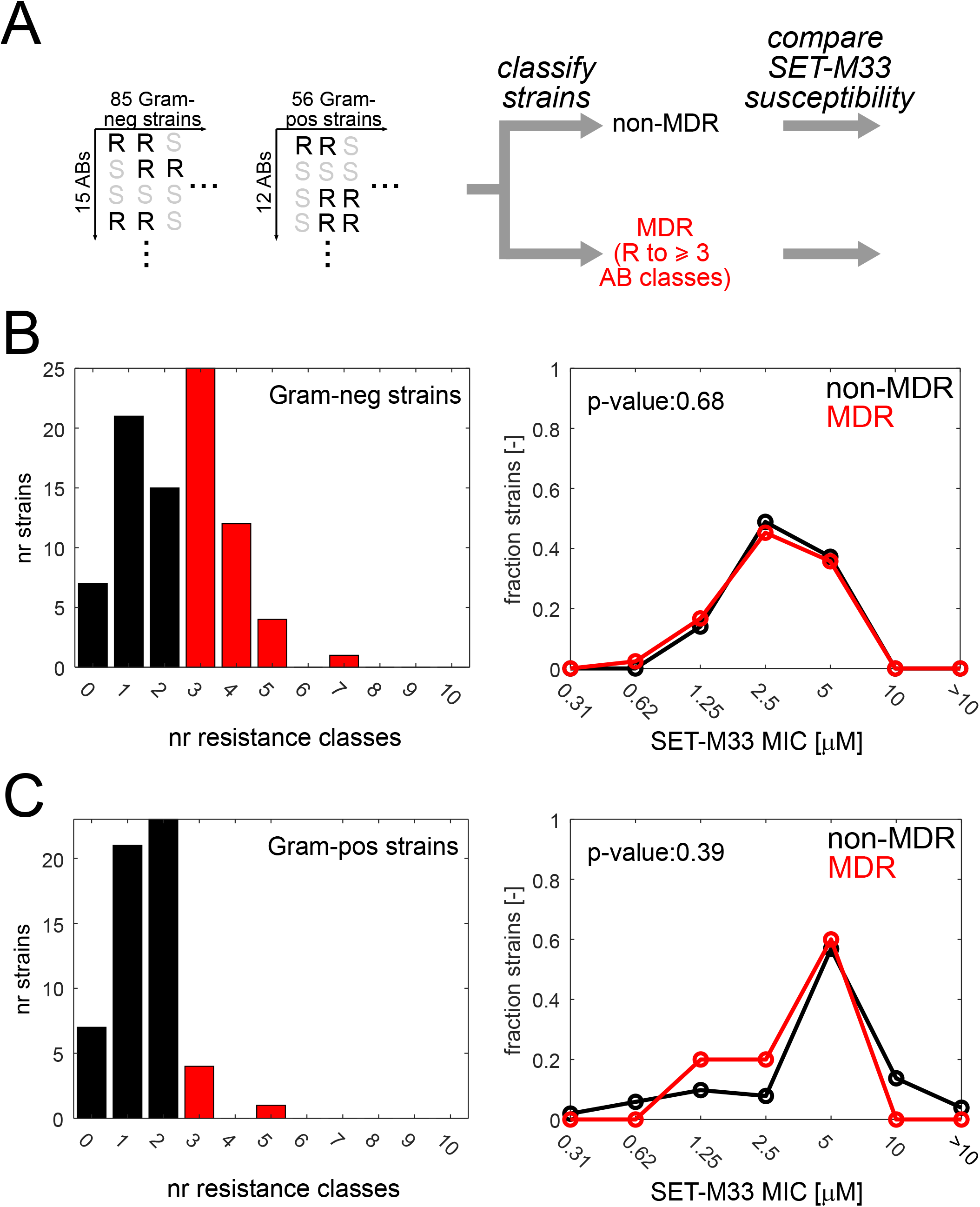
Relationship between multidrug-resistance (MDR) status and SET-M33 susceptibility in poultry isolates. **A)** Schematic of approach. **B)** Left: Distribution of antibiotic resistances (by class) across 85 Gram-negative strains. Strains resistant to 3 or more antibiotic classes are defined as MDR (red bars). Right: Distribution of SET-M33 MIC in non-MDR (black) and MDR (red) strains. P-value from a two-sample Wilcoxon ranksum test: 0.68. **C)** Left: Distribution of antibiotic resistances (by class) across 56 Gram-positive strains. Strains resistant to 3 or more antibiotic classes are defined as MDR (red bars). Right: Distribution of SET-M33 MIC in non-MDR (black) and MDR (red) strains. P-value from a two-sample Wilcoxon ranksum test: 0.39.

Moreover, there were no significant differences in SET-M33 susceptibility when classifying strains based on their resistance status to individual antibiotics (**Supplementary Figure 2**). Thus, these data suggested that the diverse resistances to conventional antibiotics carried by these strains do not affect their susceptibility to SET-M33, not even for strains that are resistant to multiple distinct antibiotic classes.

Finally, we wanted to examine the antimicrobial activity of SET-M33 against these pathogens in presence of the host. Towards this end, we established a high-throughput *in vitro* infection model that uses chicken intestinal organoid-derived monolayers to test SET-M33 efficacy in 96-well plate format (see **Figure 3A** for a schematic of the approach, and **Supplementary Text 1** and **Supplementary Figures 3**,**4** for details on the model development). Test experiments revealed that SET-M33 did cause moderate toxicity in this cultivation system, but nevertheless maintained host cell viability and epithelial barrier function at concentrations of up to 3 µM (**Supplementary Figures 3**,**4**). Having established the host-compatible range of SET-M33 concentrations, we next tested whether SET-M33 could effectively inhibit pathogen growth on organoid-pathogen co-cultures. Specifically, we focused on the Gram-negative pathogens, since all *E. faecalis* isolates had SET-M33 MIC values of 5 µM and above (outside the host-compatible range), and *E. cecorum* did not growth in the organoid cultivation medium. For each Gram-negative pathogen, we selected three isolates with SET-M33 MIC values of up to 2.5 µM (i.e. within the host-compatible concentration range) and treated each organoid-pathogen co-culture with 2-3 distinct SET-M33 concentrations (around or above the pathogen’s MIC, **Figure 3B-E**).

**Figure 3.**
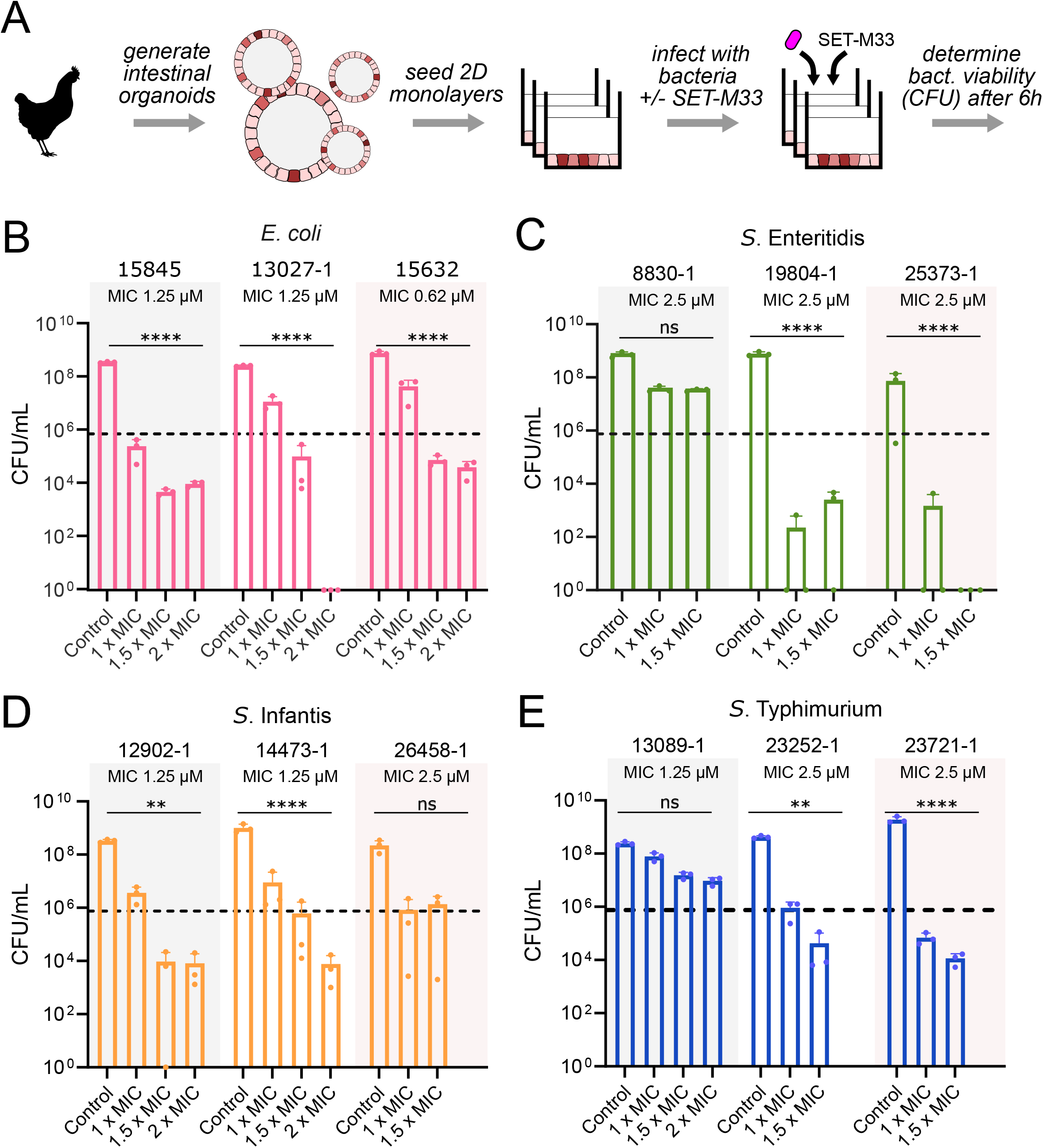
Antimicrobial activity of SET-M33 in organoid-pathogen co-cultures. **A)** Schematic of approach. **B-E)** Colony-forming units of **B)** *E. coli*, **C)**, *S*. Enteritidis, **D)** *S*. ser. Infantis, **E)** *S*. ser. Typhimurium isolates in organoid-pathogen co-cultures when treated with SET-M33 at different concentrations, compared to untreated cultures. The data represents mean ±standard deviation from one experiment with three technical replicates (****p<0.0001, **p<0.01, ns: no statistical difference of a two-way ANOVA test). Horizontal dashed lines denote the starting bacterial concentration. Small circles denote individual replicates.

We found that the antimicrobial activity of SET-M33 in these organoid-pathogen co-cultures was not only concentration, but also strain dependent: in ten out of the twelve strains, CFU counts of SET-M33 treated cultures were either around the inoculum (suggesting complete growth inhibition), or below (suggesting bactericidal SET-M33 activity), and increasing concentrations of the peptide significantly reduced bacterial growth up to 8-log (**Figure 3B-E**). Only one strain of *S*. Enteritidis (8830-1) and one of *S*. Typhimurium (13089-1) were not impacted by SET-M33 treatment even at the highest concentration used (**Figure 4C, E**). Overall, these data indicate that SET-M33 is highly effective against diverse chicken pathogens at host-compatible concentrations.

## Discussion

Although antibiotic consumption has significantly declined in recent years in European poultry farming, AMR bacteria are frequently found, and globally poultry production systems play a significant role in the spread of AMR. Coupled with the risk of limited availability of effective antimicrobials and the rapid rise of MDR strains, this challenge calls for a multifaceted approach. One promising strategy is to invest in the development of novel antimicrobials that are less susceptible to the development of resistance and capable of overcoming existing resistance mechanisms in clinically relevant bacterial species. In this context, AMPs emerge as attractive candidates. In this case study, we examined the AMP SET-M33 as a potential antimicrobial against MDR and non-MDR strains from poultry in bacterial monocultures, as well as in organoid-microbe co-cultures.

Overall, our data suggest that SET-M33 retains its activity against MDR isolates. The absence of a clear difference in SET-M33 susceptibility between MDR and non-MDR isolates suggests that its activity is not affected by AMR genes the strains may carry. This finding is aligned with broader literature showing that AMR is not necessarily correlated with reduced susceptibility to AMPs. For example, a study in *E. coli* showed that MDR can even be associated with collateral susceptibility to AMPs rather than cross-resistance (59). Besides that, cross-resistance to AMPs is often limited and depends more on their mechanisms of action than on the resistance profile itself (60,61). Similarly to what was observed in our study, cathelicidins exhibit strong antimicrobial activity against clinical relevant bacterial species, regardless of their resistance profile (62). Likewise, AMP mimetics such as ceragenins have demonstrated similarly preserved activity against both vancomycin-susceptible and -resistant enterococci (63).

Importantly, SET-M33 also retains its activity against strains that do carry resistances against other AMPs, in our case daptomycin. Daptomycin resistance is mostly associated with alterations in cell membrane charge and composition, having effects in membrane fluidity, phospholipid composition and cell wall thickness (64). All these components may in principle affect the activity of other AMPs such as SET-M33 (64). However, although many of the *E. cecorum* strains tested here were resistant to daptomycin, they were still susceptible to SET-M33 (to a similar degree as strains that are daptomycin-susceptible, see **Supplementary Figure 2**). One caveat is that in our collection of strains, only one was resistant to colistin, an antimicrobial peptide-like molecule (cyclic lipopeptide) commonly used in human medicine. As for daptomycin, recent literature has suggested that resistance to colistin can be associated with AMP cross-resistance (64,65). Mechanistic studies have shown that colistin resistance is mediated through a decrease in the net negative charge along the cell surface, resulting in lower affinity to the positive charge of colistin (66). This resistance mechanism is likely highly relevant for cationic AMPs as well, including SET-M33. Nevertheless, the single colistin-resistant isolate in our collection, a *S*. Typhimurium strain, had a similar SET-M33 MIC (5 µM) as the colistin-susceptible *S*. Typhimurium strains (median of 2.5 µM), although further studies will be needed to explore the relationship between colistin-resistance and SET-M33 susceptibility more deeply.

Despite the high effectiveness of SET-M33 against these pathogens, we also observed that SET-M33 has a toxic effect on host cells (represented here as chicken intestinal organoid cultures) at high *in vitro* doses, which constrains its usable concentration range. Previous *in vitro* studies reported cytotoxic effects of SET-M33 in bronchial epithelial monolayers with an EC_50_- of approximately 20 µM (67). In contrast, in the organoid-based system developed in this work, SET-M33 exhibited cytotoxicity at substantially lower concentrations (EC_50_-≈4 µM). The higher cytotoxicity on our organoid-based cultivation system suggests that simplified *in vitro* models based on single or immortalized cell types may underestimate the cytotoxic potential of AMPs. The increased sensitivity observed in organoids likely reflects their higher physiological complexity, including the presence of multiple cell types and more representative cell–cell interactions (68). One caveat is that here we tested chicken intestinal organoids from a single animal, and it is conceivable that the cytotoxicity of SET-M33 may vary across individuals (i.e. individuals from different chicken breeds). Nevertheless, these findings highlight the relevance of testing AMPs in a representative environment of the final targeted tissue which contributes to a more comprehensive assessment of AMP performance.

This study has several limitations. First, the resistance profile of the tested strains was only evaluated phenotypically, and a deeper understanding of the molecular mechanisms underlying these phenotypes would bring more clarity to how SET-M33 maintains its activity even against strains that are resistant to other AMPs. In line with this, evaluation of the killing kinetics would help elucidate the extent and effectiveness of SET-M33 activity against both MDR and non-MDR bacteria.

Second, this study focused on direct single exposure of SET-M33 against avian field strains, without assessing the effects of prolonged exposure. Future approaches could include evaluating potential MIC shifts after extended exposure periods as a means of detecting acquired resistance. Moreover, if resistance to SET-M33 emerges in certain strains, it would be valuable to investigate possible cross-resistance to other peptides and conventional antibiotics.

Third, the assessment of host response was primarily restricted to cytotoxicity and epithelial barrier integrity. However, AMPs may exert more subtle effects on host tissues, including alterations in cellular composition, differentiation state, or functional responses. Efforts towards tackling this issue are currently hampered by a lack of well-established protocols in most animal species. For example, there are currently no established protocols for the immunofluorescence-based detection of cell types in chicken intestinal organoid cultures. Future efforts may use advanced approaches such as single-cell transcriptomics to provide deeper insight into the impact of SET-M33 on chicken intestinal tissue composition, particularly at sub-cytotoxic concentrations.

In conclusion, this study shows that SET-M33 is highly effective *in vitro* against diverse intestinal pathogens isolated from poultry. This efficacy extends not only to MDR isolates and isolates that are resistant to other AMPs, but also to organoid-pathogen co-cultures at host-compatible concentrations, supporting its further investigation as a novel antimicrobial agent.

## Supporting information

Data set 1

## Abbreviations

AMR: antimicrobial resistance
AMP: antimicrobial peptide
APEC: avian pathogenic *E. coli*
MDR: multi drug-resistance
MIC: minimal inhibitory concentration

## Data availability

AMR profiles and SET-M33 MICs for all tested strains are included as Data set 1.

## Acknowledgements

We thank all veterinarians, clinicians, and technicians in the poultry sector for their trust in CESAC, as well as the CESAC laboratory staff for their excellent work. This project received support from a European Union Horizon Europe MSCA DN-ID grant (101073263, “Stop Spread Bad Bugs”consortium). AP acknowledges funding from the Italian Ministry of Education, University and Research (PRIN 2022, CUP B53D23003680006). GC and KK acknowledge support by the Spanish Ministry of Research and Innovation (RYC2021-033035-I /AEI/10.13039/501100011033 to KK, PLEC2022-009171 /AEI/10.13039/501100011033 to KK and GC). CERCA Programme from the Generalitat de Catalunya is also acknowledged.

## Author contributions

Conceived and designed the study: ALPL, AM, MCC, KK. Performed experiments and analyses: ALPL, AM, TA, NA, NN, KK. Supervised experiments and analyses: GC, MCC, KK. Provided material: NAN, MB, CF, AP. Wrote manuscript with contributions from all authors: ALPL, AM, MCC, KK.

## References

1. Murray CJL, Ikuta KS, Sharara F, Swetschinski L, Robles Aguilar G, Gray A, et al. Global burden of bacterial antimicrobial resistance in 2019: a systematic analysis. The Lancet. 2022 Feb;399(10325):629–55. doi:10.1016/S0140-6736(21)02724-0

2. Medina E, Pieper Helmut D. Tackling threats and future problems of multidrug-resistant bacteria. In: Stadler M, Dersch P, editors. How to Overcome the Antibiotic Crisis: Facts, Challenges, Technologies and Future Perspectives [Internet]. Cham: Springer International Publishing;2016 [cited 2025 Jun 17]. (Current Topics in Microbiology and Immunology). Available from: https://link.springer.com/10.1007/978-3-319-49284-1 doi:10.1007/978-3-319-49284-1

3. Altevogt BM, Taylor P, Akwar HT, Graham DW, Ogilvie LA, Duffy E, et al. A One Health framework for global and local stewardship across the antimicrobial lifecycle. Commun Med. 2025 Oct 7;5(1):414. doi:10.1038/s43856-025-01090-4

4. One Health Joint Plan of Action, 2022–2026 [Internet]. FAO;UNEP;WHO;WOAH;2022. Available from: http://www.fao.org/documents/card/en/c/cc2289en doi:10.4060/cc2289en

5. Li S, Wang K, Wang D, Wang H, Zhao H, Pu J, et al. Distribution and environmental dissemination of antibiotic resistance genes in poultry farms and surrounding ecosystems. Poultry Science. 2025 Jan;104(1):104665. doi:10.1016/j.psj.2024.104665

6. De Farias BO, Saggioro EM, Montenegro KS, Magaldi M, Santos HSO, Gonçalves-Brito AS, et al. Metagenomic insights into plasmid-mediated antimicrobial resistance in poultry slaughterhouse wastewater: antibiotics occurrence and genetic markers. Environmental Science and Pollution Research. 2024 Oct 12;31(51):60880–94. doi:10.1007/s11356-024-35287-2

7. Napit R, Gurung A, Poudel A, Chaudhary A, Manandhar P, Sharma AN, et al. Metagenomic analysis of human, animal, and environmental samples identifies potential emerging pathogens, profiles antibiotic resistance genes, and reveals horizontal gene transfer dynamics. Scientific Reports. 2025 Apr 9;15(1):12156. doi:10.1038/s41598-025-90777-8

8. Abreu R, Semedo-Lemsaddek T, Cunha E, Tavares L, Oliveira M. Antimicrobial drug resistance in poultry production: Current status and innovative strategies for bacterial control. Microorganisms. 2023 Apr 6;11(4):953. doi:10.3390/microorganisms11040953

9. Sweileh WM. Global research activity on antimicrobial resistance in food-producing animals. Archives of Public Health. 2021 Dec;79(1):49. doi:10.1186/s13690-021-00572-w

10. Regulation (EC) No 1831/2003 of the European Parliament and of the Council of 22 September 2003 on additives for use in animal nutrition [Internet]. Regulation (EC) 1831/2003. Available from: http://data.europa.eu/eli/reg/2003/1831/oj

11. Regulation (EU) 2019_ of the European Parliament and of the Council of 11 December 2018 on veterinary medicinal products and repealing Directive 2001_82_EC [Internet]. Regulation (EU) 2019/6. Available from: https://eur-lex.europa.eu/eli/reg/2019/6/oj/eng

12. Luiken REC, Van Gompel L, Munk P, Sarrazin S, Joosten P, Dorado-García A, et al. Associations between antimicrobial use and the faecal resistome on broiler farms from nine European countries. Journal of Antimicrobial Chemotherapy. 2019 Sep 1;74(9):2596–604. doi:10.1093/jac/dkz235

13. Ferreira M, Leão C, Clemente L, Albuquerque T, Amaro A. Antibiotic susceptibility profiles and resistance mechanisms to β-Lactams and polymyxins of *Escherichia coli* from broilers raised under intensive and extensive production systems. Microorganisms. 2022 Oct 16;10(10):2044. doi:10.3390/microorganisms10102044

14. Cerqueira-Cézar CK, Sampaio ANDCE, Caron EFF, Dellaqua TT, Ribeiro LFM, Tadielo LE, et al. Antimicrobial resistance in chicken meat: Comparing *Salmonella, Escherichia coli*, and *Enterococcus* from conventional and antibiotic-free productions. Microorganisms. 2025 Sep 23;13(10):2227. doi:10.3390/microorganisms13102227

15. Vanacker M, Lenuzza N, Rasigade JP. The fitness cost of horizontally transferred and mutational antimicrobial resistance in *Escherichia coli*. Frontiers in Microbiology. 2023 Jun 30;14:1186920. doi:10.3389/fmicb.2023.1186920

16. Magana M, Pushpanathan M, Santos AL, Leanse L, Fernandez M, Ioannidis A, et al. The value of antimicrobial peptides in the age of resistance. The Lancet Infectious Diseases. 2020 Sep;20(9):e216–30. doi:10.1016/S1473-3099(20)30327-3

17. Baindara P, Kumari S, Dinata R, Mandal SM. Antimicrobial peptides: evolving soldiers in the battle against drug-resistant superbugs. Molecular Biology Reports. 2025 Dec;52(1):432. doi:10.1007/s11033-025-10533-z

18. Flachbartova Z, Pulzova L, Bencurova E, Potocnakova L, Comor L, Bednarikova Z, et al. Inhibition of multidrug resistant *Listeria monocytogenes* by peptides isolated from combinatorial phage display libraries. Microbiological Research. 2016 Jul;188–189:34–41. doi:10.1016/j.micres.2016.04.010

19. Ramada MHS, Brand GD, Abrão FY, Oliveira M, Filho JLC, Galbieri R, et al. Encrypted antimicrobial peptides from plant proteins. Scientific Reports. 2017 Oct 16;7(1):13263. doi:10.1038/s41598-017-13685-6

20. Mohamed MF, Abdelkhalek A, Seleem MN. Evaluation of short synthetic antimicrobial peptides for treatment of drug-resistant and intracellular *Staphylococcus aureus*. Scientific Reports. 2016 Jul 11;6(1):29707. doi:10.1038/srep29707

21. Cai X, Orsi M, Capecchi A, Köhler T, Van Delden C, Javor S, et al. An intrinsically disordered antimicrobial peptide dendrimer from stereorandomized virtual screening. Cell Reports Physical Science. 2022 Dec;3(12):101161. doi:10.1016/j.xcrp.2022.101161

22. Miller RD, Iinishi A, Modaresi SM, Yoo BK, Curtis TD, Lariviere PJ, et al. Computational identification of a systemic antibiotic for Gram-negative bacteria. Nature Microbiology. 2022 Sep 26;7(10):1661–72. doi:10.1038/s41564-022-01227-4

23. Gomes B, Augusto MT, Felício MR, Hollmann A, Franco OL, Gonçalves S, et al. Designing improved active peptides for therapeutic approaches against infectious diseases. Biotechnology Advances. 2018 Mar;36(2):415–29. doi:10.1016/j.biotechadv.2018.01.004

24. Wan F, Wong F, Collins JJ, De La Fuente-Nunez C. Machine learning for antimicrobial peptide identification and design. Nature Reviews Bioengineering. 2024 Feb 26;2(5):392–407. doi:10.1038/s44222-024-00152-x

25. Torres MDT, Pedron CN, Higashikuni Y, Kramer RM, Cardoso MH, Oshiro KGN, et al. Structure-function-guided exploration of the antimicrobial peptide polybia-CP identifies activity determinants and generates synthetic therapeutic candidates. Commun Biol. 2018 Dec 7;1(1):221. doi:10.1038/s42003-018-0224-2

26. De La Fuente-Nunez C. Toward autonomous antibiotic discovery. mSystems. 2019 Jun 25;4(3):e00151–19. doi:10.1128/mSystems.00151-19

27. Zhang C, Yang M. Antimicrobial peptides: from design to clinical application. Antibiotics. 2022 Mar 6;11(3):349. doi:10.3390/antibiotics11030349

28. Nagarajan D, Roy N, Kulkarni O, Nanajkar N, Datey A, Ravichandran S, et al. Omega76: A designed antimicrobial peptide to combat carbapenem- and tigecycline-resistant *Acinetobacter baumannii*. Science Advances. 2019.

29. Suchi SA, Nam KB, Kim YK, Tarek H, Yoo JC. A novel antimicrobial peptide YS12 isolated from *Bacillus velezensis* CBSYS12 exerts anti-biofilm properties against drug-resistant bacteria. Bioprocess and Biosystems Engineering. 2023 Jun;46(6):813–28. doi:10.1007/s00449-023-02864-7

30. Chen S, Qi H, Zhu X, Liu T, Fan Y, Su Q, et al. Screening and identification of antimicrobial peptides from the gut microbiome of cockroach *Blattella germanica*. Microbiome. 2024 Dec 21;12(1):272. doi:10.1186/s40168-024-01985-9

31. Torres Salazar BO, Dema T, Schilling NA, Janek D, Bornikoel J, Berscheid A, et al. Commensal production of a broad-spectrum and short-lived antimicrobial peptide polyene eliminates nasal *Staphylococcus aureus*. Nature Microbiology. 2023 Dec 18;9(1):200–13. doi:10.1038/s41564-023-01544-2

32. Heselpoth RD, Euler CW, Fischetti VA. PaP1, a broad-spectrum lysin-derived cationic peptide to treat polymicrobial skin infections. Frontiers in Microbiology. 2022 Mar 10;13:817228. doi:10.3389/fmicb.2022.817228

33. Yu G, Baeder DY, Regoes RR, Rolff J. Predicting drug resistance evolution: insights from antimicrobial peptides and antibiotics. Proceedings of the Royal Society. 2018.

34. Kintses B, Méhi O, Ari E, Számel M, Györkei Á, Jangir PK, et al. Phylogenetic barriers to horizontal transfer of antimicrobial peptide resistance genes in the human gut microbiota. Nature Microbiology. 2018 Dec 17;4(3):447–58. doi:10.1038/s41564-018-0313-5

35. Lewis K. The science of antibiotic discovery. Cell. 2020 Apr;181(1):29–45. doi:10.1016/j.cell.2020.02.056

36. Li T, Wang Z, Guo J, De La Fuente-Nunez C, Wang J, Han B, et al. Bacterial resistance to antibacterial agents: Mechanisms, control strategies, and implications for global health. Science of The Total Environment. 2023 Feb;860:160461. doi:10.1016/j.scitotenv.2022.160461

37. Nuri R, Shprung T, Shai Y. Defensive remodeling: How bacterial surface properties and biofilm formation promote resistance to antimicrobial peptides. Biochimica et Biophysica Acta (BBA) - Biomembranes. 2015 Nov;1848(11):3089–100. doi:10.1016/j.bbamem.2015.05.022

38. EFSA Panel on Animal Health and Welfare (AHAW), Nielsen SS, Bicout DJ, Calistri P, Canali E, Drewe JA, et al. Assessment of animal diseases caused by bacteria resistant to antimicrobials: Poultry. EFS2. 2021 Dec;19(12). doi:10.2903/j.efsa.2021.7114

39. Nolan LK, Vaillancourt J, Barbieri NL, Logue CM. Colibacillosis. In: Swayne DE, Boulianne M, Logue CM, McDougald LR, Nair V, Suarez DL, et al., editors. Diseases of Poultry [Internet]. 1st ed. Wiley;2020 [cited 2026 Mar 20]. p. 770–830. Available from: https://onlinelibrary.wiley.com/doi/10.1002/9781119371199.ch18 doi:10.1002/9781119371199.ch18

40. Mak PHW, Rehman MA, Kiarie EG, Topp E, Diarra MS. Production systems and important antimicrobial resistant-pathogenic bacteria in poultry: a review. Journal of Animal Science and Biotechnology. 2022 Dec 14;13(1):148. doi:10.1186/s40104-022-00786-0

41. Gao J, Duan X, Li X, Cao H, Wang Y, Zheng SJ. Emerging of a highly pathogenic and multidrug resistant strain of *Escherichia coli* causing an outbreak of colibacillosis in chickens. Infection, Genetics and Evolution. 2018 Nov;65:392–8. doi:10.1016/j.meegid.2018.08.026

42. Gast RK, Porter RE. *Salmonella* Infections. In: Swayne DE, Boulianne M, Logue CM, McDougald LR, Nair V, Suarez DL, et al., editors. Diseases of Poultry [Internet]. 1st ed. Wiley;2020 [cited 2026 Mar 20]. p. 717–53. Available from: https://onlinelibrary.wiley.com/doi/10.1002/9781119371199.ch16 doi:10.1002/9781119371199.ch16

43. Sharma S, Kaur S, Naguib M, Bragg A, Schneider A, Kulkarni RR, et al. Major foodborne bacterial pathogens in poultry: implications for human health and the poultry industry and probiotic mitigation strategies. Microorganisms. 2025 Oct 14;13(10):2363. doi:10.3390/microorganisms13102363

44. Jung A, Chen LR, Suyemoto MM, Barnes HJ, Borst LB. A review of Enterococcus cecorum infection in poultry. Avian Diseases. 2018 Sep;62(3):261–71. doi:10.1637/11825-030618-Review.1

45. Arias CA, Murray BE. The rise of the *Enterococcus*: beyond vancomycin resistance. Nature Reviews Microbiology. 2012 Apr;10(4):266–78. doi:10.1038/nrmicro2761

46. Rehman MA, Yin X, Zaheer R, Goji N, Amoako KK, McAllister T, et al. Genotypes and phenotypes of Enterococci isolated from broiler chickens. Frontiers in Sustainable Food Systems. 2018 Dec 13;2:83. doi:10.3389/fsufs.2018.00083

47. Pini A, Falciani C, Mantengoli E, Bindi S, Brunetti J, Iozzi S, et al. A novel tetrabranched antimicrobial peptide that neutralizes bacterial lipopolysaccharide and prevents septic shock *in vivo*. The FASEB Journal. 2010 Apr;24(4):1015–22. doi:10.1096/fj.09-145474

48. Van Der Weide H, Vermeulen-de Jongh DMC, Van Der Meijden A, Boers SA, Kreft D, Ten Kate MT, et al. Antimicrobial activity of two novel antimicrobial peptides AA139 and SET-M33 against clinically and genotypically diverse *Klebsiella pneumoniae* isolates with differing antibiotic resistance profiles. International Journal of Antimicrobial Agents. 2019 Aug;54(2):159–66. doi:10.1016/j.ijantimicag.2019.05.019

49. Van Der Weide H, Brunetti J, Pini A, Bracci L, Ambrosini C, Lupetti P, et al. Investigations into the killing activity of an antimicrobial peptide active against extensively antibiotic-resistant *K. pneumoniae* and *P. aeruginosa*. Biochimica et Biophysica Acta (BBA) - Biomembranes. 2017 Oct;1859(10):1796–804. doi:10.1016/j.bbamem.2017.06.001

50. Brunetti J, Falciani C, Roscia G, Pollini S, Bindi S, Scali S, et al. *In vitro* and *in vivo* efficacy, toxicity, bio-distribution and resistance selection of a novel antibacterial drug candidate. Scientific Reports. 2016 May 12;6(1):26077. doi:10.1038/srep26077

51. Johnson TJ, Wannemuehler Y, Doetkott C, Johnson SJ, Rosenberger SC, Nolan LK. Identification of minimal predictors of avian pathogenic *Escherichia coli* virulence for use as a rapid diagnostic tool. Journal of Clinical Microbiology. 2008 Dec;46(12):3987–96. doi:10.1128/JCM.00816-08

52. Laurentie J, Mourand G, Grippon P, Furlan S, Chauvin C, Jouy E, et al. Determination of epidemiological cutoff values for antimicrobial resistance of *Enterococcus cecorum*. Barrs VR, editor. Journal of Clinical Microbiology. 2023 Mar 23;61(3):e01445–22. doi:10.1128/jcm.01445-22

53. Magiorakos AP, Srinivasan A, Carey RB, Carmeli Y, Falagas ME, Giske CG, et al. Multidrug-resistant, extensively drug-resistant and pandrug-resistant bacteria: an international expert proposal for interim standard definitions for acquired resistance. Clinical Microbiology and Infection. 2012 Mar;18(3):268–81. doi:10.1111/j.1469-0691.2011.03570.x

54. Lourenço ALP, Rouam El Khatab O, Falciani C, Pini A, Aragon V, Cerdà-Cuéllar M, et al. SET-M33 peptide as a selective *in vitro* antimicrobial agent against the porcine respiratory pathogen *Glaesserella parasuis*. Zhao H, editor. Microbiology Spectrum. 2026 Apr 7;14(4):e03918–25. doi:10.1128/spectrum.03918-25

55. Brunetti J, Carnicelli V, Ponzi A, Di Giulio A, Lizzi AR, Cristiano L, et al. Antibacterial and anti-inflammatory activity of an antimicrobial aeptide synthesized with D amino acids. Antibiotics. 2020 Nov 24;9(12):840. doi:10.3390/antibiotics9120840

56. Cresti L, Falciani C, Cappello G, Brunetti J, Vailati S, Melloni E, et al. Safety evaluations of a synthetic antimicrobial peptide administered intravenously in rats and dogs. Scientific Reports. 2022 Nov 11;12(1):19294. doi:10.1038/s41598-022-23841-2

57. EFSA, ECDC. European Union summary report on antimicrobial resistance in zoonotic and indicator bacteria from animals and food in the European Union in 2009. EFS2. 2011 Jul;9(7). doi:10.2903/j.efsa.2011.2154

58. Falciani C, Lozzi L, Pollini S, Luca V, Carnicelli V, Brunetti J, et al. Isomerization of an antimicrobial peptide broadens antimicrobial spectrum to Gram-positive bacterial pathogens. Bereswill S, editor. PLoS ONE. 2012 Oct 2;7(10):e46259. doi:10.1371/journal.pone.0046259

59. Lázár V, Martins A, Spohn R, Daruka L, Grézal G, Fekete G, et al. Antibiotic-resistant bacteria show widespread collateral sensitivity to antimicrobial peptides. Nature Microbiology. 2018 May 24;3(6):718–31. doi:10.1038/s41564-018-0164-0

60. Spohn R, Daruka L, Lázár V, Martins A, Vidovics F, Grézal G, et al. Integrated evolutionary analysis reveals antimicrobial peptides with limited resistance. Nature Communications. 2019 Oct 4;10(1):4538. doi:10.1038/s41467-019-12364-6

61. Kintses B, Jangir PK, Fekete G, Számel M, Méhi O, Spohn R, et al. Chemical-genetic profiling reveals limited cross-resistance between antimicrobial peptides with different modes of action. Nature Communications. 2019 Dec 16;10(1):5731. doi:10.1038/s41467-019-13618-z

62. Veldhuizen EJA, Brouwer EC, Schneider VAF, Fluit AC. Chicken cathelicidins display antimicrobial activity against multiresistant bacteria without inducing strong resistance. Cloeckaert A, editor. PLoS ONE. 2013 Apr 22;8(4):e61964. doi:10.1371/journal.pone.0061964

63. Hacioglu M, Yilmaz FN, Oyardi O, Bozkurt Guzel C, Inan N, Savage PB, et al. Antimicrobial activity of ceragenins against vancomycin-susceptible and -resistant *Enterococcus* spp. Pharmaceuticals. 2023 Nov 23;16(12):1643. doi:10.3390/ph16121643

64. Mishra NN, McKinnell J, Yeaman MR, Rubio A, Nast CC, Chen L, et al. *In Vitro* cross-resistance to daptomycin and host defense cationic antimicrobial peptides in clinical methicillin-resistant *Staphylococcus aureus* isolates. Antimicrobial Agents and Chemotherapy. 2011 Sep;55(9):4012–8. doi:10.1128/AAC.00223-11

65. Jangir PK, Ogunlana L, Szili P, Czikkely M, Shaw LP, Stevens EJ, et al. The evolution of colistin resistance increases bacterial resistance to host antimicrobial peptides and virulence. eLife. 2023 Apr 25;12:e84395. doi:10.7554/eLife.84395

66. Mousavi SMJ, Hosseinpour M, Kodori M, Rafiei F, Mahmoudi M, Shahraki H, et al. Colistin antibacterial activity, clinical effectiveness, and mechanisms of intrinsic and acquired resistance. Microbial Pathogenesis. 2025 Apr;201:107317. doi:10.1016/j.micpath.2025.107317

67. Cresti L, Conte G, Cappello G, Brunetti J, Falciani C, Bracci L, et al. Inhalable polymeric nanoparticles for pulmonary delivery of antimicrobial peptide SET-M33: Antibacterial activity and toxicity *in vitro* and *in vivo*. Pharmaceutics. 2022 Dec 20;15(1):3. doi:10.3390/pharmaceutics15010003

68. Zhao D, Farnell MB, Kogut MH, Genovese KJ, Chapkin RS, Davidson LA, et al. From crypts to enteroids: establishment and characterization of avian intestinal organoids. Poultry Science. 2022 Mar;101(3):101642. doi:10.1016/j.psj.2021.101642

