## Supplementary Information for "*In vitro* efficacy of synthetic antimicrobial peptide SET-M33 against poultry isolates with diverse antimicrobial resistance phenotypes"

**Supplementary text 1 – developing experimental protocol for testing SET-M33 in a chicken intestinal organoid model**

The use of organoid models for antimicrobial peptide testing requires culture conditions that preserve both organoid viability and peptide stability. To address this technical challenge, we selected different culture media, specifically organoid growth medium (OGM), representing a standard medium for organoid cultures, DMEM, DMEM + FBS and ADV DMEM/F-12. We evaluated the effect of each medium on the stability of two SET-M33 isomers, namely SET-M33L (version used in the rest of the manuscript) and SET-M33D (stereoisomer only used during the development of this experimental protocol).

First, we tested SET-M33L and SET-M33D stability indirectly by incubating the peptides in each medium for different time intervals (1, 2, 4, 8 and 24 h), hereafter referred as “conditioning time”. Following incubation, we tested the antimicrobial activity of the conditioned peptide solutions against the *E. coli* strain ATCC25922, which is highly susceptible to SET-M33 (**Supplementary Figure 3A**). The results showed how OGM markedly affected the antimicrobial activity of both isomers. In particular, peptide activity was significantly reduced after 2 h in the medium (**Supplementary Figure 3B**). Across all conditioning times, *E. coli* growth levels were comparable to those observed in the control (medium without peptide), indicating that the components of OGM impair the activity of SET-M33. A similar trend was observed in DMEM + FBS, where peptide activity was also impaired, particularly for SET-M33D (**Supplementary Figure 3C**). In this case, loss of antimicrobial activity was already evident after 1 h of conditioning. Moreover, SET-M33L showed a highly variable activity in DMEM + FBS. In contrast, peptide activity was preserved in DMEM and ADV DMEM/F-12. In both media, the two SET-M33 isomers retained the ability to significantly reduce *E. coli* growth compared with the control (**Supplementary Figure 3D, E**), although only ADV DMEM/F-12 fully inhibited bacterial growth across all conditioning times (**Supplementary Figure 3E**).

Second, we assessed organoid viability in the same selection of media. Organoids were grown in 2D monolayers and once they reached confluency, the standard cultivation medium (i.e. OGM) was exchanged for the tested media, and viability was measured after 24h of incubation through metabolic assay (**Supplementary Figure 3F**). Organoids cultured with DMEM and DMEM + FBS showed a substantial reduction in viability (40%) compared with the standard cultivation media OGM, while the metabolic activity of the cells remained comparable when ADV DMEM/F-12 was used (**Supplementary Figure 3E**). These results indicate that ADV DMEM/F-12 preserves both SET-M33 antimicrobial activity and organoid viability over 24 h. We therefore selected this medium for all subsequent experiments.

**Supplementary tables**

**Supplementary table 1.**

| **Antibiotic** | **Abbreviation** | **Class** | **Mode of action** |
| --- | --- | --- | --- |
| Ampicillin | AMP | Penicillins | Cell wall synthesis inhibitor |
| Meropenem | MERO | Carbapenems | Cell wall synthesis inhibitor |
| Ceftazidime | TAZ | Cephalosporins | Cell wall synthesis inhibitor |
| Cefotaxime | FOT | Cephalosporins | Cell wall synthesis inhibitor |
| Ciprofloxacin | CIP | Quinolones | DNA gyrase inhibition |
| Nalidixic Acid | NAL | Quinolones | DNA gyrase inhibition |
| Azithromycin | AZI | Macrolides | 50S ribosome inhibition |
| Amikacin | AMI | Aminoglycosides | 30S ribosome inhibition |
| Gentamicin | GEN | Aminoglycosides | 30S ribosome inhibition |
| Tigecycline | TGC | Glycylcyclines | 30S ribosome inhibition |
| Chloramphenicol | CHL | Phenicols | 50S ribosome inhibition |
| Colistin | COL | Polymyxins | Cell membrane disruption |
| Tetracycline | TET | Tetracyclines | 30S ribosome inhibition |
| Trimethoprim | TMP | Sulfonamides | Dihydrofolate synthesis (DHFS) inhibition |
| Sulfamethoxazole | SMX | Sulfonamides | Dihydropteroate synthesis (DHPS) inhibition |
| Vancomycin | VAN | Glycopeptides | Cell wall inhibition |
| Teicoplanin | TEI | Glycopeptides | Cell wall inhibition |
| Quinupristin/dalfopristin | SYN | Streptogramins | 50S ribosome inhibition |
| Daptomycin | DAP | Cyclic lipopeptides | Membrane depolarization |
| Erythromycin | ERY | Macrolides | 50S ribosome inhibition |
| Linezolid | LZD | Oxazolidinones | 50S ribosome inhibition |

**Supplementary Figures**

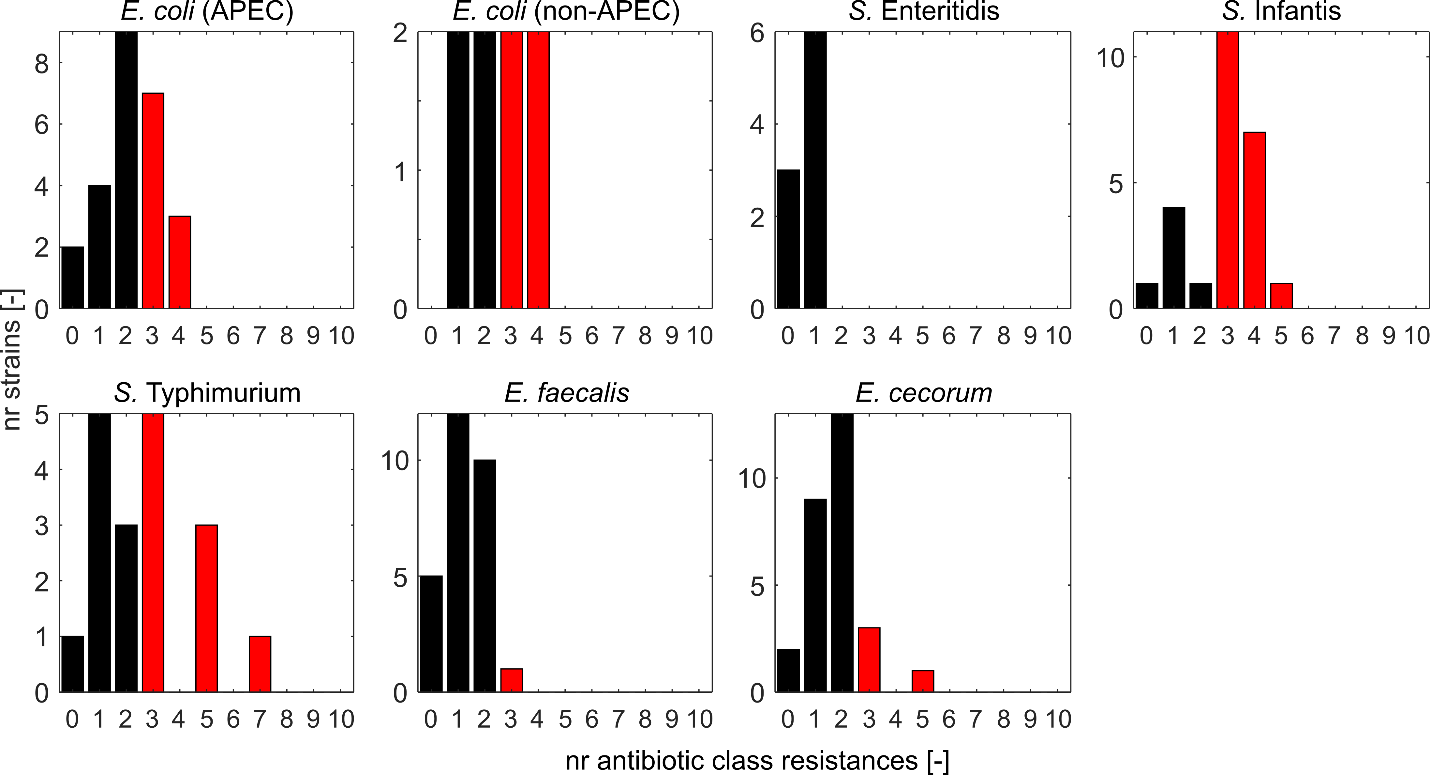

**Supplementary Figure 1. Distribution of the number of antibiotic resistances (by class) separated by strain type.** Strains resistant to 3 or more antibiotic classes are defined as multidrug-resistant (red bars). APEC, avian pathogenic *E. coli*.

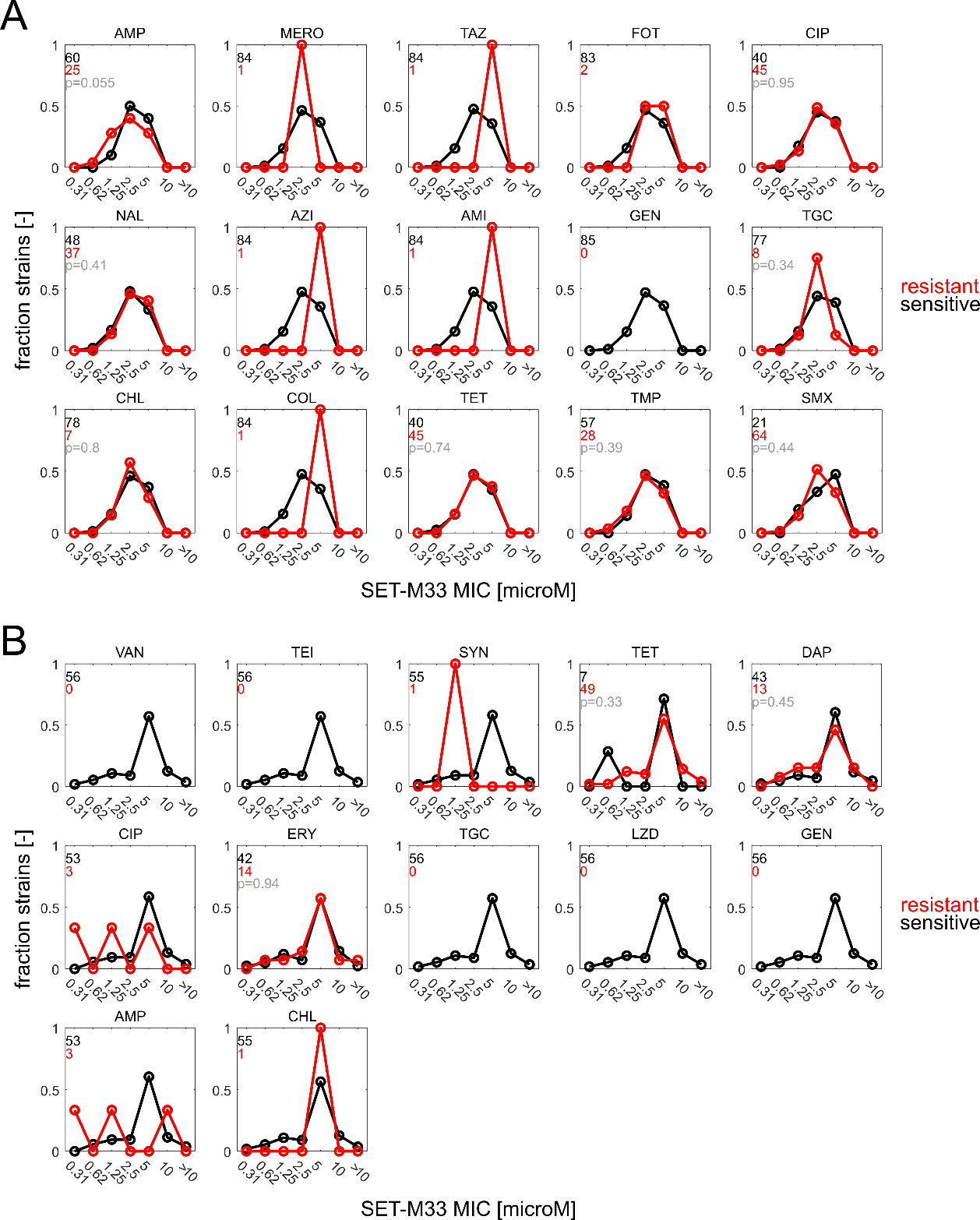

**Supplementary Figure 2. Distribution of SET-M33 MIC in Gram-negative (A) and Gram-positive (B) isolates, separated by antibiotic resistance state.** Black: strains sensitive to respective antibiotic. Red: strains resistant to respective antibiotic. P-values: result of two-sided Wilcoxon rank sum test (shown in gray, and only calculated for cases where at least 5 strains are resistant to the respective antibiotic). Antibiotics: ampicillin (AMP), meropenem (MERO), ceftazidime (TAZ), cefotaxime (FOT), ciprofloxacin (CIP), nalidixic acid (NAL), azithromycin (AZI), amikacin (AMI), gentamicin (GEN), tigecycline (TGC), chloramphenicol (CHL), colistin (COL), tetracycline (TET), trimethoprim (TMP), sulfamethoxazole (SMX), vancomycin (VAN), teicoplanin (TEI), quinupristin/dalfopristin (SYN), daptomycin (DAP), erythromycin (ERY), linezolid (LZN).

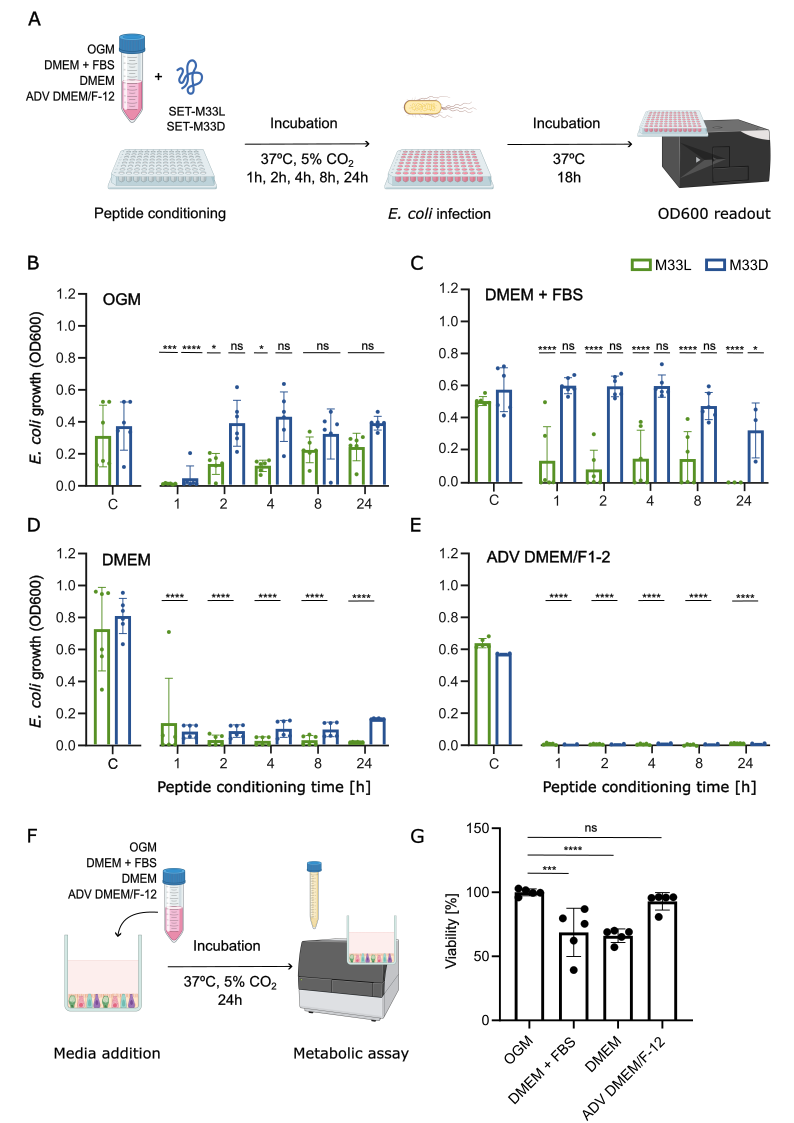
**Supplementary Figure 3. SET-M33 stability assessment and organoids viability in different media.** **(A)** Experimental workflow used to evaluate the effect of different media on SET-M33 stability. Antimicrobial activity against *E. coli* ATCC25922 was quantified following conditioning in **(B)** OGM, **(C)** DMEM + 20% FBS, **(D)** DMEM, and **(E)** ADV DMEM/F-12. **(F)** Workflow used to evaluate organoid viability after exposure to each medium. **(G)** Organoid viability measured by metabolic assay after 24 h incubation. The data were obtained from two independent experiments with three technical replicates per experiment and are shown as mean ± standard deviation (**** p<0.0001, *** p < 0.001, * p<0.05, ns: no statistical difference, of a two-way ANOVA test). Circles denote the individual replicates.

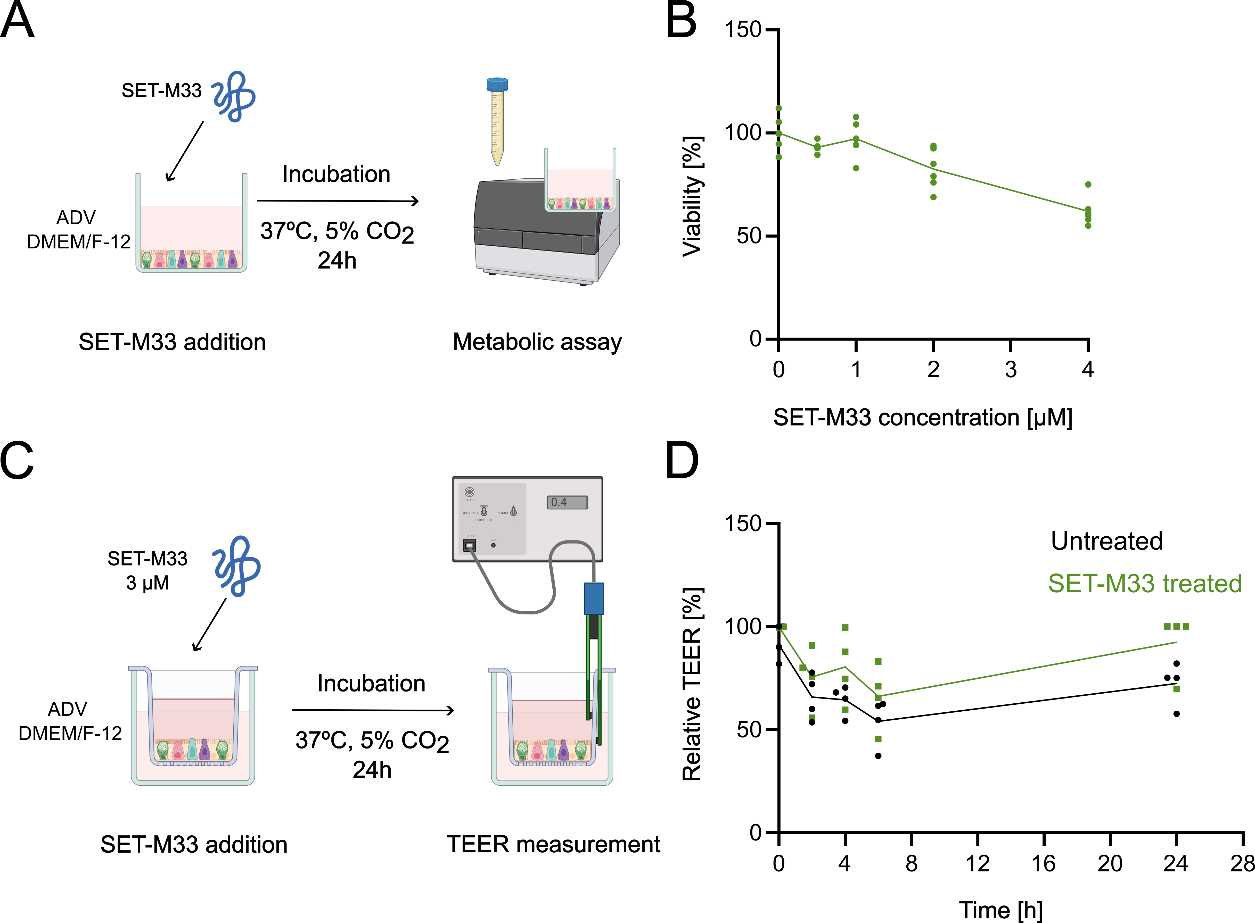

**Supplementary Figure 4. Cytotoxicity of SET-M33 in chicken intestinal organoid-derived epithelial monolayers and epithelial barrier integrity. (A)** Schematic of the methods used to test SET-M33 cytotoxicity. **(B)** Viability of chicken intestinal organoids when treated with increasing concentration of SET-M33. Two independent experiments with three technical replicates were performed. **(C)** Methodology employed to investigate the impact of SET-M33 on organoid tissue. **(D)** Trans-epithelial electrical resistance (TEER) time course of the organoid-derived 2D monolayer cultures seeded in transwell when treated with 3 µM of SET-M33 (The TEER absolute value at T0 accounted for 110.72 ± 34.46 Ωxcm^2^). The data was generated from two independent experiments with two technical replicates per experiment. The replicates are shown as individual points and all the data are presented as mean ± standard deviation.
